# Comparative phylogenomic analyses of SNP versus full locus datasets: insights and recommendations for researchers

**DOI:** 10.1101/2023.09.02.556036

**Authors:** Jacob S. Suissa, Gisel Y. De La Cerda, Leland C. Graber, Chloe Jelley, David Wickell, Heather R. Phillips, Ayress D. Grinage, Corrie S. Moreau, Chelsea D. Specht, Jeff J. Doyle, Jacob B. Landis

## Abstract

**Premise:** In phylogenomic analyses, no consensus exists on whether using single nucleotide polymorphisms (SNPs) or including flanking regions (full ‘locus’) is best, nor how strictly missing data should be filtered. Moreover, empirical evidence on whether SNP-only trees are suitable for downstream phylogenetic comparative methods such as divergence time estimation and ancestral state reconstructions is lacking.

**Methods:** Using GBS data from 22 taxa of *Glycine*, we addressed the effects of SNP vs. locus usage and filtering stringency on phylogenomic inference and phylogenetic comparative methods. We compared branch length, node support, and divergence time estimation across eight datasets with varying amounts of missing data and total size.

**Results:** Our results reveal five aspects of phylogenomic data usage: **1**. tree topology is largely congruent regardless of data type or filtering parameters; **2**. filtering missing data too strictly reduces the confidence in some relationships; **3**. absolute branch lengths vary by two orders of magnitude between datasets; **4**. data type and branch length variation have little effect on divergence time estimation; **5**. phylograms significantly alter the estimation of ancestral states.

**Discussion:** When conducting phylogenomic analyses we recommend not to filter datasets too strictly to minimize the risk of misleading topologies, low support, and inaccurate divergence times.

## Premise of the study

With increased efficiency in high throughput sequencing, generating genome-wide data for phylogenomic analysis is cheaper and more feasible (Young and Gillung, 2020). There are many different approaches that can be used to generate sequence data for phylogenetic questions—each come with their own limitations and necessary inputs (Chambers et al., 2023). Both wet lab (e.g., access to living tissue, difficulty in DNA extractions, and laboratory costs) and computational requirements (e.g., reference genome, initial bioinformatic investment such as probe design, and ultimate bioinformatic investment such as SNP calling) need to be considered before starting a phylogenomics project (McKain et al., 2018; Dodsworth et al., 2019).

While many phylogenomic methods rely on gene sequences, these data are not easily generated in some systems and approaches (Dodsworth et al., 2019). Single nucleotide polymorphisms (SNPs) are useful in phylogenomic studies in part due to the ease in data collection (i.e., genome skimming, RAD-Seq, or genome resequencing) and their broad distribution across the genome (Leaché and Oaks, 2017), and their phylogenetic information. However, the effects of SNP vs. locus datasets on tree topology, branch lengths, and nodal support are uncertain and the applicability of using inferred trees from SNPs in downstream phylogenetic comparative methods (PCMs) has been called into question (Leaché et al., 2015).

While there is no shortage of work discussing recommendations for Hyb-Seq phylogenomic studies (Weitemier et al., 2014; Kadlec et al., 2017; McKain et al., 2018; Villaverde et al., 2018) or using a SNP-based method (often generated with RAD-Seq or other genotyping by sequencing methods; Lee et al., 2014; Hyun et al., 2020, 2021), there is little overlap in studies comparing results from different approaches. One such comparison by Zhou and Xiang (2022) demonstrated that congruence was found in topology and divergence time dating in their SNP vs locus datasets. However, conflict may exist based on how the SNP data is handled in analyses. For example, topologies and node support can differ if the SNP and surrounding flanking regions are treated as genes/loci in coalescent approaches, including Astral (Mirarab et al., 2014). This may lead to considerable incongruence primarily due to the lack of signal in the gene tree from each locus, as usually inferred by either RAxML (Stamatakis, 2014) or IQ-TREE (Minh et al., 2020). However, in coalescent approaches such as SVDquartets (Chifman and Kubatko, 2014), each nucleotide is taken individually so the invariant flanking regions are not necessary since these sites would be phylogenetically uninformative in regards to relationships. Therefore, further evaluation of how these different kinds of datasets impact phylogenomic inferences is necessary, especially when downstream PCMs are used.

Another longstanding issue in phylogenomic inference is the effect of missing data. Given the complexity and scale of genome-wide sequencing projects, it is almost certain that datasets will contain some degree of missing information, whether that be from biological reasons or limitations in the sequencing approach. This raises the important question of how missing data affect the accuracy and reliability of phylogenetic inference on a genomic scale. In some cases, missing data has not been demonstrated to have a direct impact on phylogenetic inference, especially when datasets are large and the amount of missing data is relatively low (Wiens, 2003, 2006; Philippe et al., 2004). For instance, Weins (2003) and Roure et al., (2013) showed that even with moderate amounts of missing data, the resulting phylogenetic trees were still highly congruent with those obtained from complete datasets. A remarkable example of this is the successful resolution of a recent radiation event in Acanthaceae using RAD-Seq data with as much as 90% missing data (Tripp et al., 2017). These findings suggest that the impact of missing data on phylogenetic inference might be mitigated when taxonomic datasets are sufficiently large.

However, conflicting viewpoints exist regarding the influence of missing data on phylogenetic inference. Some argue that missing data can have a negative impact by exacerbating uncertainty in inferred topologies, potentially leading to the estimation of misleading branch lengths (Huelsenbeck, 1991; Lemmon et al., 2009). One such study found that 70% data completeness was necessary to avoid spurious relationships (Smith et al., 2020). These studies contend that missing data can introduce biases, affecting the accuracy and reliability of the inferred phylogenetic relationships. Consequently, these uncertainties may propagate further in downstream analyses, potentially compromising the interpretation of evolutionary processes. Considering these contrasting results, the effect of missing data on phylogenetic inference, especially with genome-wide data, remains a topic of considerable debate and best practices are currently unknown.

To explore the effects of SNP vs. locus data usage, and the stringency for when a SNP is kept or removed by filtering, on phylogenomic inference and PCMs, we used the soybean genus *Glycine* Willd. (Fabaceae; Ratnaparkhe et al., 2011). Members of *Glycine* subgenus *Glycine* are perennial species with diverse habits ranging from vines to subshrubs, native to Australia and occurring on various Pacific islands (Hermann, 1962). There are approximately 26 named perennial species (Sherman-Broyles et al., 2014b; Landis and Doyle, 2023), with many taxa still informally recognized (Gunner and Doyle, 2014). Species of *Glycine* were initially placed into genome groups based on infertility of artificial crosses (Singh and Hymowitz, 1985; Hymowitz et al., 1998), which have largely been supported with molecular phylogenies and now are the basis for genome group classifications (Sherman-Broyles et al., 2014a). Taxonomic issues remain in the subgenus, particularly involving the *G. tomentella* and *G. tabacina* complexes, both of which include allopolyploids and the diploid taxa that hybridized to produce them ∼0.5 million years ago (mya; Bombarely et al., 2014; Sherman-Broyles et al., 2014b, 2017). In this study, we use GBS data from 22 diploid *Glycine* species to investigate the effects of data usage and filtering stringency for common phylogenetic inference and downstream PCMs. Specifically, we set out to compare the effects of SNPs vs. entire RAD loci on overall inferred tree topology, node support, branch length, age, and downstream phylogenetic comparative methods. We aim to provide support for researchers in deciding the best phylogenetic approach in their study system when using genome-wide data.

## Methods

### SNP vs. locus identification

The GBS data used in this study was previously described in Landis and Doyle (2023). Briefly, DNA was extracted from tissue grown from seed stocks from CSIRO (Commonwealth Scientific and Industrial Research Organization, Australia) using a cetyltrimethylammonium bromide (CTAB) method (Doyle and Doyle, 1987). Extracted DNA was sent to the University of Wisconsin Biotechnology Center for GBS library preparation following (Elshire et al., 2011) using the *ApeK1* restriction enzyme. Libraries were sequenced on an Illumina HiSeq2000 at Cornell University Institute of Biotechnology with a single-end 100 bp approach. Reads were demultiplexed using the process_radtags perl script in Stacks v2.55 (Rochette et al., 2019). Accession numbers for the Sequence Read Archive (SRA), number of reads, and which genome group each accession belongs to can be found in Table 1. Since a *de novo* SNP calling approach was undertaken instead of a reference guided approach, raw reads were cleaned using fastp v0.12.4 (Chen et al., 2018) by automatically detecting adapters and requiring a minimum quality score of 20 and a minimum read length of 60 bp.

**Table 1.** Accessions used in this study, including number of reads, estimated depth after SNP calling, number of loci assembled within each accession, and the known genome group.

| Species | SRA number | Number of reads | Estimated depth | Assembled loci | Genome group |
| --- | --- | --- | --- | --- | --- |
| *Glycine albicans* | SRR18315602 | 3,319,398 | 18.92x | 177,436 | I-genome |
| *Glycine arenaria* | SRR18315601 | 1,416,973 | 16.20x | 88,094 | H-genome |
| *Glycine argyrea* | SRR18315591 | 3,133,554 | 22.07x | 143,284 | A-genome |
| *Glycine canescens* | SRR18315580 | 2,409,732 | 17.95x | 135,312 | A-genome |
| *Glycine clandestina* | SRR18315575 | 5,317,539 | 32.37x | 166,063 | A-genome |
| *Glycine sp. "cracens"* | SRR18315595 | 3,204,690 | 21.30x | 152,150 | B-genome |
| *Glycine falcata* | SRR18315574 | 687,974 | 11.00x | 63,079 | F-genome |
| *Glycine gracei* | SRR18315573 | 2,707,429 | 19.52x | 139,795 | A-genome |
| *Glycine hirticaulis* | SRR18315572 | 1,491,560 | 15.15x | 99,371 | H-genome |
| *Glycine lactovirens* | SRR18315571 | 3,229,367 | 20.42x | 160,799 | I-genome |
| *Glycine latifolia* | SRR18315600 | 2,240,664 | 17.65x | 128,166 | B-genome |
| *Glycine microphylla* | SRR18315599 | 1,825,974 | 16.03x | 114,992 | B-genome |
| *Glycine pindanica* | SRR18315598 | 1,972,682 | 16.74x | 118,742 | H-genome |
| *Glycine pullenii* | SRR18315597 | 1,151,892 | 13.25x | 87,740 | H-genome |
| *Glycine stenophita* | SRR18315592 | 2,434,744 | 20.42x | 119,952 | B-genome |
| *Glycine syndetika* | SRR18315589 | 3,883,250 | 21.99x | 178,332 | A-genome |
| *Glycine tomentella D1* | SRR18315587 | 3,595,114 | 20.78x | 175,600 | E-genome |
| *Glycine tomentella D2* | SRR18315586 | 2,240,510 | 13.58x | 166,821 | E-genome |
| *Glycine tomentella D3* | SRR18315585 | 3,315,188 | 19.67x | 170,003 | D-genome |
| *Glycine tomentella D5A* | SRR18315581 | 2,861,702 | 19.53x | 148,274 | Ha-genome |
| *Glycine tomentella D5B* | SRR18315579 | 2,461,459 | 20.15x | 123,171 | H-genome |
| *Glycine sp. "wilsonii"* | SRR18315593 | 3,884,757 | 24.91x | 157,579 | B-genome |

The denovo_map.pl script in Stacks v2.62 (Rochette et al., 2019) was used to *de novo* call SNPs with default parameters with the inclusion of --force-diff-len in ustacks to account for slight variation in read lengths across samples due to different barcode sizes. Specifically for SNP identification, the default value for the minimum stack depth (-m 3) was used, meaning that at least three reads were needed to determine an allele. The distance between stacks (-M 2) was used to minimize combining repetitive regions together into one locus. The populations module was then used with different filtering criteria for the proportion of accessions that a SNP must be present in to be retained (r=0% - one individual, 15% - three individuals, 30% - six individuals, 45% - nine individuals, 60% - 13 individuals, 75% - 16 individuals, 90% - 19 individuals, and 100% - 22 individuals) plus the additional filtering parameters of minimum MAF (minor allele frequency) = 0.05, minimum MAC (minor allele count) = 3, and maximum observed heterozygosity = 0.5. For the eight datasets, variant (SNP only) and full loci (SNP + flanking regions) were exported in phylip format using the --phylip-var-all and --phylip-var commands in the populations module. Due to file recognition issues, the exported phylip files from Stacks were converted to fasta format for Bayesian analysis (see below) with AliView v1.28 (Larsson, 2014). The amount of missing data per accession in each filtered dataset was calculated in VCFtools v0.1.16 (Danecek et al., 2011) from the VCF file that was generated alongside the phylip alignments by the populations module.

### Phylogenetic inference

Maximum likelihood trees were inferred with RAxML-NG v1.1.0 (Kozlov et al., 2019) with 100 bootstraps, the GTR+G model of molecular evolution. We specified *G. falcata* as the outgroup, as demonstrated by previous nuclear studies showing *G. falcata* sister to the rest of the perennial species (Hwang et al., 2019; Zhuang et al., 2022; Landis and Doyle, 2023). No partitioning was done in the full locus dataset to make comparisons between variant and full locus as equitable as possible. Datasets with the different levels of minimum number of accessions sharing a given site (0% - one individual, 15% - three individuals, 30% - six individuals, 45% - nine individuals, 60% - 13 individuals, 75% - 16 individuals, 90% - 19 individuals, and 100% - 22 individuals), as well as variant sites only (SNPs) and all sites (variant sites plus invariant flanking regions in the full RAD locus) were used to infer maximum likelihood trees. Missing data at any given site was coded as an N since Ns, gaps, and missing data are treated the same way in RAxML (Kozlov et al., 2019).

### Divergence time estimation

Divergence time dating was done with BEAST v2.7.2 (Bouckaert et al., 2019) using a secondary calibration based on previous dating analyses of *Glycine* (Zhuang et al., 2022) incorporating the split of the *Glycine* and *Phaseolus* lineages ∼22-37 mya (Koenen et al., 2021). The following settings were used in generating the BEAUTi file: ambiguities allowed, estimate substitution rate, gamma category count of 4, estimate gamma shape, GTR substitution model with empirical base frequencies, optimized relaxed clock, birth-death tree prior. A calibration point forcing a monophyletic clade of all *Glycine* species with a median value of 6.12 and a sigma of 0.515 with a normal distribution was set. A total of 100-500 million MCMC sampling every 5000 or until the effective sample size values were over 200 as checked for convergence with Tracer v1.7.2 (Rambaut et al., 2018). An initial comparison between runs using a yule tree prior or a birth-death prior showed overlapping age estimates for all nodes observed. Therefore, all resulting analyses were done with the birth-death model.

### Comparison of filtering and data type on branch lengths and nodal support

Branch lengths and node support were compared between data types (SNP vs. locus) and filtering stringency. These analyses were only conducted on RAxML trees since BEAST tree branch lengths and node ages were all standardized to unit time. To compare branch lengths between locus and SNP trees across different filtering parameters, all RAxML trees were read into R using the read.tree function in ape v5.7-1 (Paradis and Schliep, 2019). Branch lengths, node support, node number, filtering stringency, and data type were extracted from each tree and aggregated into a dataframe using a custom script in R v4.2.2 (R Core team, 2021). Boxplots of branch lengths and node support were compared within data type across filtering stringencies using the geom_boxplot function in ggplot2 v3.4.2 (Wickham, 2016), where the box represents the middle 50% of the data, with the bottom and top edges of the box representing the first and third quartiles, respectively. The line inside the box represents the median. The whiskers extend from the edges of the box to the smallest and largest observations that are not outliers. Outliers, which are observations that are more than 1.5 times the interquartile range away from the nearest quartile, were plotted as individual points outside the whiskers. Pairwise comparisons of branch lengths and nodal support at different filtering levels within datasets were made using a Wilcoxon signed rank test with a Bonferroni correction with continuity correction using the pairwise.wilcox.test function in R.

### Dating phylograms

Due to the large number of taxa incorporated into phylogenetic trees in the genomics era, rate smoothing approaches are often used to convert the best tree from a maximum likelihood analysis or maximum clade credibility Bayesian framework to an ultrametric tree (Ho and Duchêne, 2014; Barba-Montoya et al., 2021; Costa et al., 2022). RelTime, which is based on the relative rate framework (RRF) (Tamura et al., 2012, 2018) and implemented in MEGA11 (Tamura et al., 2021), was used for divergence time estimation with the sequence alignments and RAxML trees as input. Not only can RelTime be a thousand times faster than Bayesian methods (Tamura et al., 2012), but RelTime estimates have been shown to be consistently more accurate than other ML methods such as TreePL and least squares dating (Barba-Montoya et al., 2021). To investigate the impacts of SNP vs. loci datasets, as well as the impact of overall missing data in an alignment, on downstream divergence time estimate we dated each of the fifteen phylogenies inferred in RAxML with the exception of the largest alignment file consisting of 111 million sites due to computational limitations. We used an equivalent calibration point to that in the BEAST analyses, setting the age of the most recent common ancestor of *Glycine* to a mean of 6.12 mya, with a standard deviation of 0.435 to span the confidence interval of 5.27-6.97 mya. A normal distribution for this calibration point was used, along with the GTR model of molecular evolution with a gamma rate. Differences between inferred nodes were compared at the same data filtering level (missing data) between variant only and locus data. A paired Student’s t-test (Student, 1908) was used to identify significant differences in inferred ages, with Bonferroni correction for multiple tests.

### Node age by data type and filtering stringency

To explore the relationship between node age and filtering stringency, we selected the mean node age for a set of selected nodes across the phylogeny for each filtered dataset. Using ggplot2 in R, we plotted mean node age and standard error across each of the selected nodes. We also conducted linear regressions to test for significant trends between node age and filtering stringency, using the lm function in R.

### Ancestral character estimation

A downstream approach that is a fundamental tool of comparative phylogenetic methods is ancestral character state estimation (Felsenstein, 1985). Different approaches for this have been reviewed in other studies, but generally all are conducted after a tree is inferred (Holland et al., 2020). The preferred approaches are to use chronograms where branch lengths are in units of time; however, one study suggested that similar results will be generated with a phylogram where branch lengths are proportional to substitutions per site (Litsios and Salamin, 2012). To further test this on trees inferred from both SNP and full locus, which have largely different branch lengths (see below) we used stochastic character mapping with simulated binary data on the phylograms and chronograms inferred using the 45% shared locus datasets to see if any differences were observable in number of transitions, time spent in different states, and percentage of traits across all nodes. Specifically, a binary trait was randomly simulated across the tips and stochastic mapping was performed in phytools v1.5-1 (Revell, 2012) with an ARD model and 1000 simulations.

We also explored how tree and data type affect ancestral character state estimations of continuous variables. To do this we first simulated a continuous trait under a Brownian motion process (assuming σ^2^ to be 1) along the SNP BEAST tree derived from the 45% shared dataset using the fastBM function in phytools v1.5-1. After simulating these continuous data, we then estimated ancestral character states across the same four selected trees for the discrete traits mentioned above using the anc.mL function in phytools v1.5-1. These include RAxML phylogram and BEAST chronogram built using SNPs and full loci with the 45% locus dataset. Observed and reconstructed values of the simulated continuous trait were projected onto the edges of the respective phylogenies using the function contMap in phytools v1.5-1.

## Results

### SNPs vs. locus information

Based on the required threshold of the percentage of accessions that a SNP must be found to be retained, from zero percent (unique SNPs in any given accession) to 100% (shared across accessions), the number of loci varied from 1,960 (100%) to 1,339,951 (0%) (Table 2). Across all loci, the number of variant sites ranged from 2,491 to 118,925; when the full locus was taken into account the total alignment size ranged from 182,544 base pairs (bp) to 111,994,953 bp. The mean fraction of missing data for just the variant sites across all individuals in a filtering threshold ranged from zero to 0.522; while the mean level of missing data in the full locus ranged from 0.099 to 0.906. Missing data in any given accession across datasets is shown in Supplemental Table S1, showing that most accessions were similar in terms of amount of missing data, except for *G. falcata* which showed substantially more missing sites than other accessions.

**Table 2.** Summary of the different datasets including fraction of shared sites a SNPs must have to be retained, number of loci for each filtering scheme, the number of base pairs for variant and locus datasets, as well as the median and mean values of missing data for each dataset.

| Shared percentage required | Loci | Variant sites (bp) | All sites (bp) | median missing variant | mean missing variant | median missing locus | mean missing locus |
| --- | --- | --- | --- | --- | --- | --- | --- |
| 100 | 1,960 | 2,491 | 182,544 | 0.000 | 0.000 | 0.115 | 0.099 |
| 90 | 6,522 | 9,125 | 606,786 | 0.033 | 0.050 | 0.133 | 0.140 |
| 75 | 13,305 | 19,956 | 1,236,145 | 0.092 | 0.121 | 0.178 | 0.200 |
| 60 | 22,713 | 33,340 | 2,102,501 | 0.162 | 0.201 | 0.248 | 0.273 |
| 45 | 43,490 | 58,081 | 3,983,546 | 0.273 | 0.322 | 0.359 | 0.392 |
| 30 | 76,196 | 85,371 | 6,906,941 | 0.378 | 0.423 | 0.483 | 0.507 |
| 15 | 170,505 | 117,310 | 15,060,214 | 0.483 | 0.518 | 0.649 | 0.663 |
| 0 | 1,339,951 | 118,925 | 111,994,953 | 0.489 | 0.522 | 0.904 | 0.906 |

### Topology

The inferred topologies across all sixteen datasets were largely congruent, regardless of whether the alignment was inferred with SNPs or the full locus with the exception of when data were filtered so that only sites shared across all accessions were retained (Fig. S1). The one part of the topology that was impacted was within the H-genome group (Fig. 1), specifically with the placement of *G. arenaria, G. hirticaulis, G. pullenii*, and *G. tomentella* D5B. In both 100% trees, *G. pullenii* is sister to the clade of *G. pindanica* and *G. tomentella* D5B with no support (29% bootstrap support in the entire locus tree and 49% bootstrap support in the variant only tree). In the other inferred trees, *G. pullenii* is sister to the rest of the H-genome clade with full support or *G. tomentella* D5B is sister to the rest of the H-genome with full support. Overall, the level of missing data, from zero to 90% did not have large impacts on the inferred topology. The five genome groups represented by multiple individuals were all recovered as monophyletic. The D and E-genome groups were sister to each other, which were in turn sister to a clade of H and Ha genome groups. This resulting clade was successively sister to the I-genome, A-genome, and B-genome group. The F-genome group represented by *G. falcata* was sister to all other perennial accessions.

**Figure 1.**
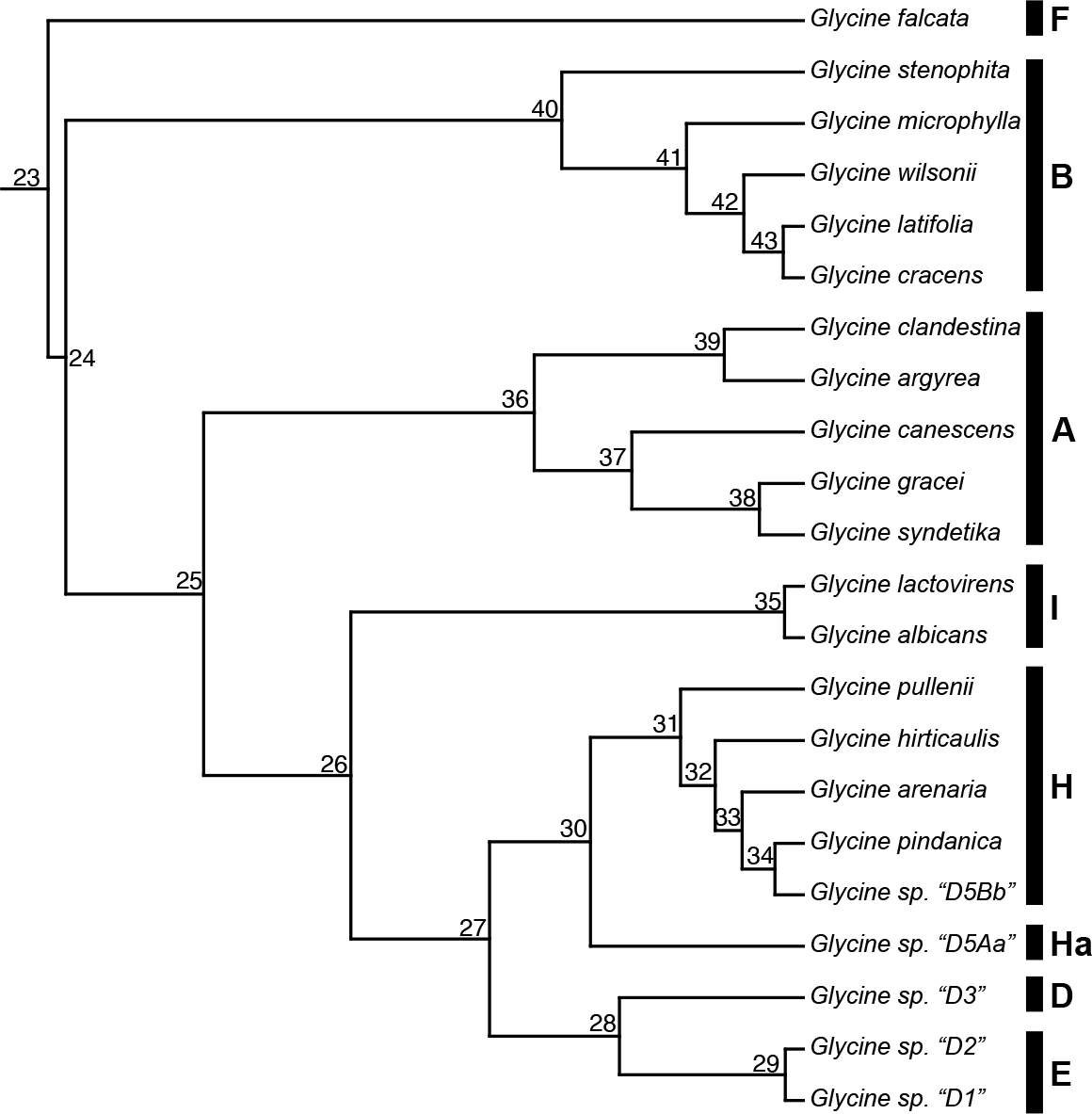
Chronogram of the SNP phylogeny, which is largely congruent across different data types and filtering parameters. The only observable differences are when SNP filtering is too extreme and only sites shared across all taxa are kept. The bars next to the tips represent the assigned genome group for each species, with the node labels being node numbers for downstream analyses. Numbers along branches denote node order (not support values). Node numbers are the same as those plotted in figures 3 and 4.

### Comparison of filtering and data type on branch lengths and node support

Branch lengths and node support were only compared within data type (i.e., between filtering stages in SNP and locus). While data on branch lengths and node support were not statistically compared between data types, one noticeable difference is that raw branch lengths of locus data were in general two orders of magnitude shorter than those of SNP data (Fig. 2). Based on Shapiro-Wilk’s test of normality (*P* value < 0.01) the branch lengths for the locus and SNP dataset were not normally distributed (Fig. 2). However, across the locus dataset, the lowest branch lengths occurred at 0% and 100% shared data, and the largest occurred between 30%– 75% shared data. Alternatively, within the SNP data branch lengths were longest in the dataset with 0% shared data and shortest in the dataset with 100% shared data (Fig. 2). Based on a Wilcoxon signed rank test with a Bonferroni correction, node support was not significantly different across filtering stringencies either within the SNP or locus data (Supplemental Table S2). However, there were differences in branch lengths across filtering stringencies (Fig. 2). Within the SNP data, significant differences in branch lengths based on a Wilcoxon signed rank test occurred between filtering stringency 0% & 75%; 0% & 100%; 15% & 30%; and 15% & 100%. Within locus data, significant differences in branch lengths based on a Wilcox signed rank test occurred between filtering stringency 0% & 15%; 0% & 30%; 0% & 45%; 0% & 60%; 0% & 75%; 0% & 100%; 15% & 60%; 15% & 75%; and 60% & 100% (Supplemental Table S2; *P* value <0.05 with Bonferroni correction).

**Figure 2.**
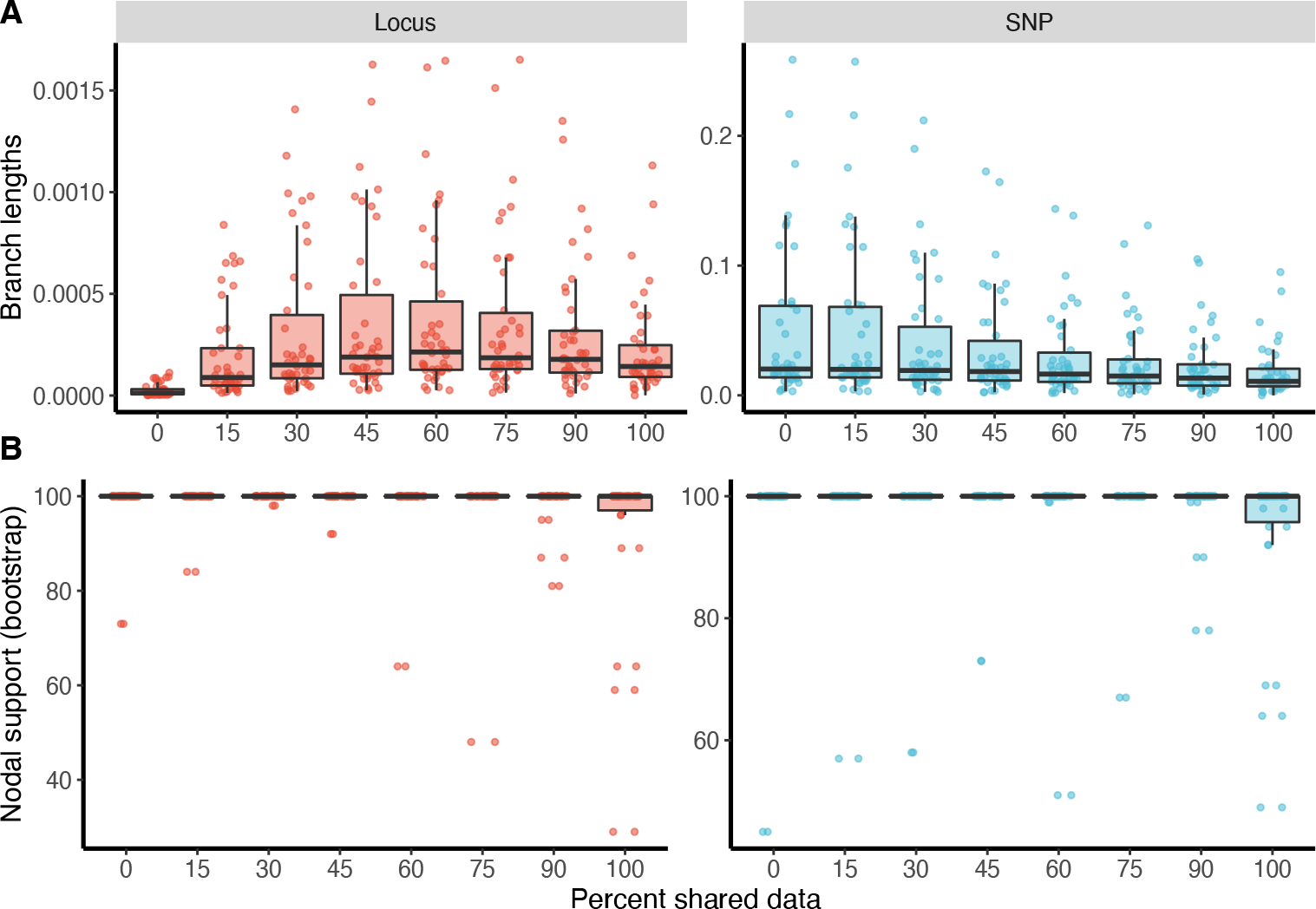
A) comparison of branch lengths between SNP and locus datasets by filtering stringency B) comparison of nodal support between SNP and locus datasets by filtering stringency.

### Divergence time estimation

The BEAST analyses within data type produced congruent results in terms of estimated node age across filtering stringency, with the 95% confidence interval for selected nodes being almost entirely overlapping (Figure 3 and 4). The one exception to this was node 34, which is in the H-genome group and is the most recent common ancestor of *G. pindanica* and *G. sp*. “D5Bb”. As noted above, the relationships within the H-genome group change slightly based on filtering stringency, with some relationships no longer being well supported. For some nodes, a significant correlation exists that when SNP filtering stringency increases, the inferred divergence time increases; in other nodes the opposite is true (Figure 3B and 4B). Even though these changes are significant, the differences are still within the 95% confidence interval of all inferred ages for that particular node across the different levels of filtering.

**Figure 3.**
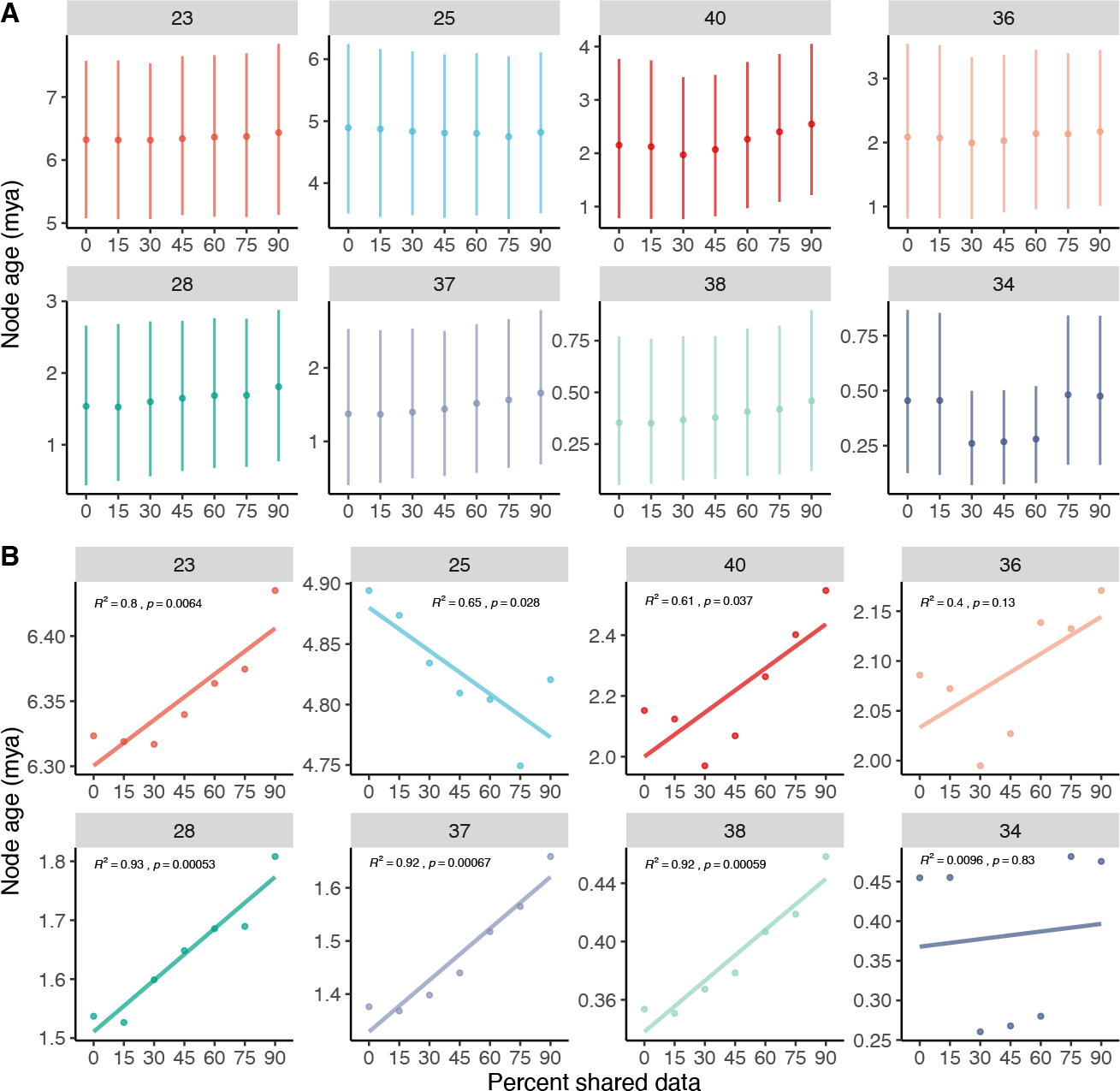
Selected nodes from BEAST node ages for SNP inferred phylogenies. A) mean node ages (point) with 95% confidence intervals of absolute age (lines). B) linear regressions depicting the relationship between mean node age and filtering stringency. Numbers above each inset represent a node number corresponding to Figure 1.

**Figure 4.**
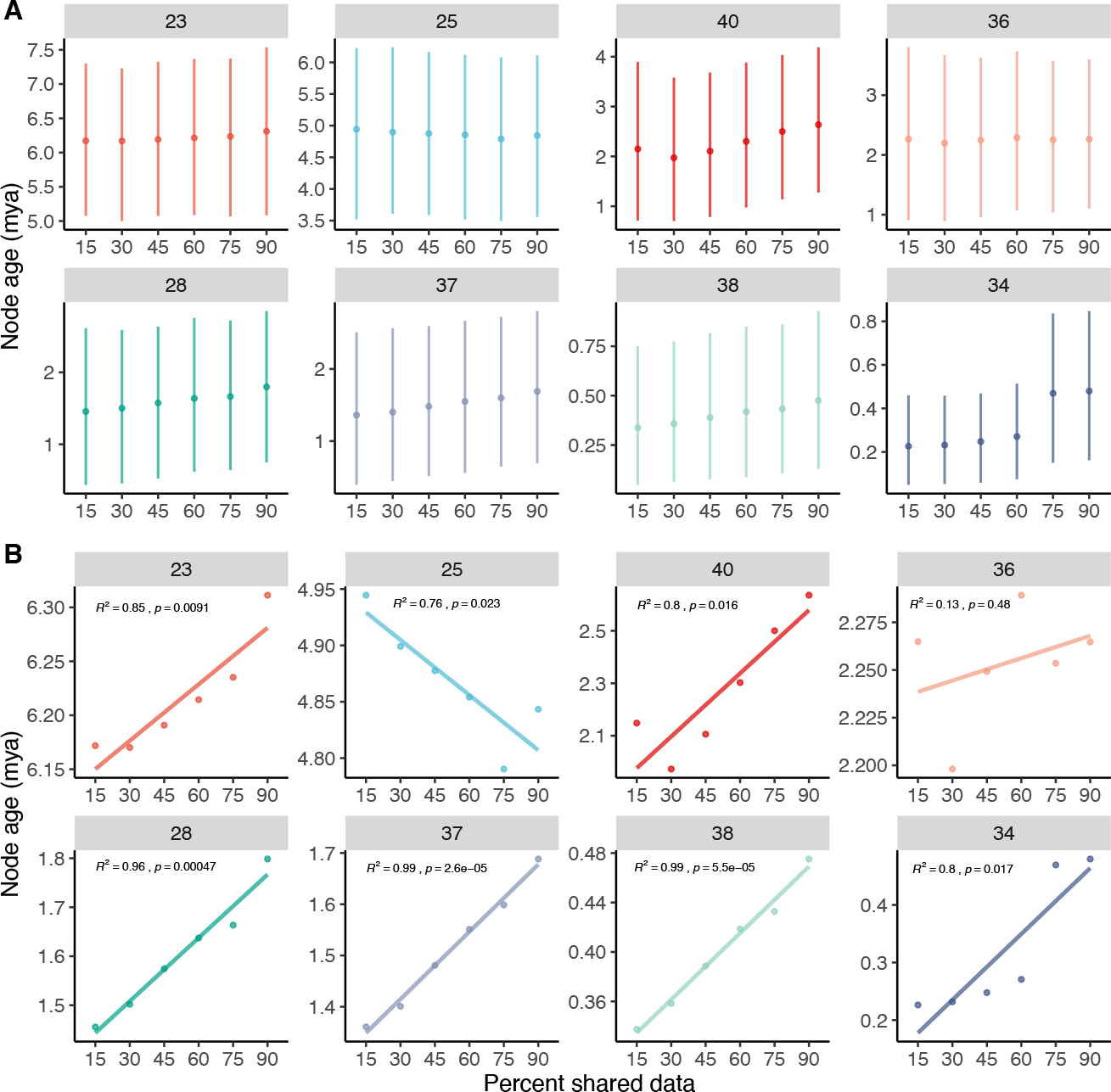
Selected nodes from BEAST node ages for full locus inferred phylogenies. A) mean node ages (point) with 95% confidence intervals of absolute age (lines). B) linear regressions depicting the relationship between mean node age and filtering stringency. Numbers above each inset represent a node number corresponding to Figure 1.

The inferred divergence times using RelTime were highly similar regardless of whether SNP or locus were used. The inferred node ages of RelTime were within the 95% confidence interval generated in the BEAST analyses. Generally speaking, as missing data increased the mean difference between variant and locus increased marginally. The mean difference in ages of the same node was 0.023 mya when no missing data was allowed (100%), increasing to 0.025 mya (90%), 0.036 mya (75%), 0.041 mya (60%), 0.038 mya (45%), 0.052 mya (30%), and 0.61 mya (15%). The differences between datasets was much less than the standard deviation of 0.435 mya used in the calibration point for dating (6.12 ± 0.435 mya). A paired Student’s t-test showed no significant difference in node ages between S NP and locus after Bonferonni correction (critical value 0.05/7); while two comparisons showed a significant differences in estimated node age at the 45% (*P* value = 0.021) and 30% (*P* value = 0.014) given a standard cutoff of 0.05.

### Ancestral character state estimation

Ancestral character state estimation differed qualitatively and quantitatively between RAxML and BEAST trees. The RAxML SNP tree had a total of 47 transitions, spending 64% of time in state B and 36% of time in State A. Transition rates were 25.556 changes per unit branch length from A–B and 41.287 changes per unit branch length from B–A. The RAxML locus tree had a total of 7 transitions, spending 83% of time in state B and 17% of time in State A. Transition rates were equal between State A and B (0.472 changes per unit branch length). The BEAST SNP tree had a total of 134 transitions, spending 65% of time in state B and 35% of time in State A. Transition rates were 2.352 changes per unit branch length from A–B and 4.301 changes per unit branch length from B–A. The BEAST locus tree had a total of 128 transitions, spending 65% of time in state B and 35% of time in State A. Transition rates were 2.241 changes per unit branch length from A–B and 4.063 changes per unit branch length from B–A.

Qualitatively, the ancestral states and proportion of time spent in each state appears to be relatively similar between the RAxML tree using SNP data and both stochastic character maps of the BEAST trees (Fig. 5). However, quantitatively the transition rates for the RAxML SNP tree were 10.9 times higher and the total number of transitions were almost 3 times lower compared to the BEAST trees. The RAxML locus tree was most different compared to all other trees. The ancestral state estimation for almost every node was incongruent, the total number of transitions were 6.7 times less than the respective SNP tree and almost 20 times less than the BEAST trees, and the transitions rates were equal between A–B and very low. Overall, using trees with branch lengths in proportion to nucleotide substitution rate appears to provide drastically different quantitative and qualitative results compared to trees with branch lengths in proportion to absolute time.

**Figure 5.**
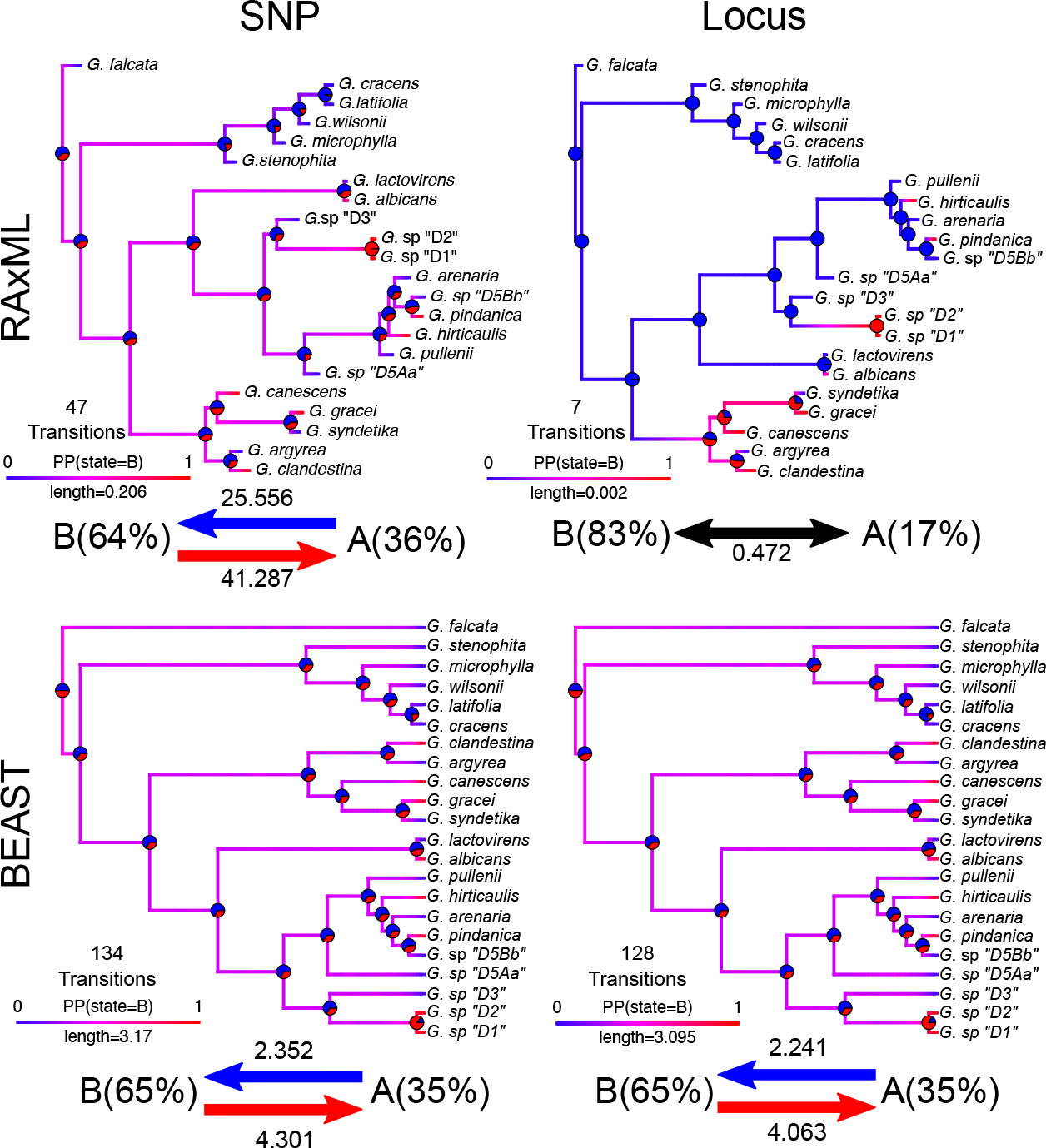
Summarized results of 1000 stochastic character maps of simulated data on two representative phylograms and chronograms from the 45% filtering stringency. Blue and red arrows represent transition rates. The percentages in parentheses are the total time spent in each state. Branches are colored by their posterior probability of being in state A or B. Pie charts represent the probability of nodes in state A or B. The total number of transitions are denoted on top of legend.

Similar disparities in ancestral state estimations were observed in continuous variables as well. Broadly, there was virtually no difference in ancestral state estimations and model parameters between SNP and locus BEAST chronograms. However, there were large differences between both SNP and locus phylograms and between RAxML phylograms and BEAST chronograms. SNP and locus-based phylograms both estimated a root ancestral state near the upper limits of the modeled continuous variable. Moreover, the rate of variance (σ^2^) was two orders of magnitude greater in the locus phylogram compared to the SNP phylogram. The root state of chronograms were estimated to be near the middle of the modeled continuous variable and the rate of variance was much lower compared to RAxML trees and was equal between SNP and locus Beast trees.

## Discussion

### Topological congruence, branch lengths, and nodal support

The topology generated here using a *de novo* SNP calling approach (Fig. 1) is congruent with the phylogeny presented in Landis and Doyle (2023), which used a reference genome approach for calling SNPs. In comparison, both the *de novo* approach presented here and the reference guided approach used in Landis and Doyle (2023) generated a fully resolved phylogeny with strong support, suggesting that robust relationships can be inferred with or without a reference genome. Moreover, the topologies we generated *de novo* using either SNP or full locus datasets were almost entirely identical. The only caveats to this were the inferred topologies from the 100% filtering, which had different relationships among members of the H-genome group, specifically *G. arenaria, G. hirticaulis*, and *G. pullenii*, than the other inferred phylogenies (Fig. S1). However, these incongruent relationships showed low (<70%) bootstrap support.

We found that the absolute branch lengths differed between full locus and SNP datasets. Branch lengths of full locus datasets were two orders of magnitude smaller than the absolute branch lengths of SNP datasets (Fig. 2). However, while the absolute branch lengths were quite different, the relative branch lengths between datasets did not appear to vary drastically. A notable observation was that patterns of filtering stringency on branch lengths differed between SNP and locus datasets. In SNP datasets, branch lengths tended to decrease with increasing filtering stringency. However, in locus datasets branch lengths were lowest with 0% shared data and largest around 45-60% shared data (Fig. 2; SI dataset). In the SNP only datasets, increased missing data led to longer branch lengths, while missing data in entire locus datasets did not show a linear pattern (Fig. 2). Even with the large differences in absolute branch lengths between the SNP and full locus datasets, the inferred node ages were largely overlapping across filtering and data types (Fig. 2). We attribute this to a similarity in relative branch lengths between the SNP and locus inferred phylogenies.

Some of the branch length discrepancies we observed might be alleviated with the incorporation of an ascertainment bias flag (Lewis, 2001; Leaché et al., 2015) as implemented in RAxML and IQ-TREE. The correction for ascertainment bias predominately impacts the calculated branch lengths, since the variant sites are sampled in a non-uniform manner and only including SNPs deviates from theoretical expectations in how bases change (Lachance and Tishkoff, 2013). However, in some cases implementing this bias correction is not straightforward since constant sites cannot be included in the alignment but the number of invariant sites do need to be counted, which for nonmodel systems and genome wide data may not be an easy endeavor. Within *Glycine*, heterozygosity is low due to a predominant selfing mode of reproduction (Doyle et al., 2004). In highly outcrossing systems, high levels of heterozygosity may also cause issues in phylogenetic inference due to the way ambiguities are handled (e.g., treating ambiguities as one base or the other, or by ignoring ambiguities). In these cases, using models of genotype evolution (eight possible genotype combinations and the rates of change between combinations) instead of models of molecular evolution (four possible bases and the rates of change between bases), as implemented in RAxML-NG (Kozlov et al., 2018), may provide a more accurate inference especially in terms of branch length estimation. One additional caveat to consider is the age of divergence between the focal species. We have shown that SNP data are useful for *Glycine*, where the perennial species diverged from the annual *G. max* approximately 8 mya (Zhuang et al., 2022). SNP data have also been successful in determining the relationships in *Drosophila*, which diverged 40-60 mya (Rubin et al., 2012). However, a SNP based approach for species that diverged over 100 mya may not provide robust results (Rubin et al., 2012). Determining the crown age of focal taxa should be considered before undertaking a SNP-based phylogenomic analysis.

### Divergence time dating by percent shared data and locus type

Previous phylogenomic studies have shown that allowing for missing data can help resolve difficult relationships and increase node support (Wiens, 1998, 2006; Wiens and Moen, 2008). However, there is a risk that allowing more missing da ta may increase long-branch attraction (Wiens, 2006), especially when the missing data are not randomly distributed among loci (Xi et al., 2016). Yet, there exists no conclusive pattern of consistent bias in branch length and missing data (Jiang et al., 2014). In our datasets, we observed a significant inverse linear relationship between filtering stringency and mean age across most nodes, where most nodes showed increased mean ages with decreased missing data (Figure 3 and 4). However, it is important to note that variation in mean age tended to be captured within the full confidence interval of the node estimation, demonstrating the importance of bounding age-dependent hypotheses by confidence intervals (Figure 3A and 4A). As such, while these correlations between filtering and mean age were significant, they may not be phylogenetically meaningful on small times scales (Bromham, 2019).

Several SNP-based studies have shown that retaining sites with missing data increase the support for phylogenetic relationships (Wagner et al., 2013; Hodel et al., 2017; Tripp et al., 2017). This is likely due to larger datasets containing more phylogenetically informative sites and because excluding sites with missing data can mask regions of the genome with the highest mutation rates. For instance, a study of African frogs (*Afrixalus* spp) showed that allowing approximately 60% missing data in a given site provided congruent tree inferences, whereas topological differences were observed between different quality of samples at 80-90% missing data (Crotti et al., 2019). Our results support the notion that the inclusion of sites with some missing data or sites where some accessions do not possess a SNP, at least to a certain extent, is beneficial in phylogenomic reconstruction (not to mention the faster computational time compared to all sites being retained). While the overall topologies generally did not change (except in the most stringent filtering criteria; Fig S1), some variation in node support across filtering stringencies was observed. In particular, the largest variation in support was found when a SNP was required to be retained in all individuals (100%) compared to all other thresholds (Figure 2). We also observed the smallest variation in node support when a SNP was required to be present in 30-45% of individuals, which is similar to best case scenarios of Crotti et al. (2019).

### Phylogenetic comparative methods and downstream analyses

We demonstrated that using phylograms leads to drastically different ancestral character estimations in both discrete and continuous characters compared to chronograms (Figure 5 and 6; also see Wilson et al., 2022). In ancestral character state estimations of both continuous and discrete states, rates of morphological evolution were higher in phylograms compared to chronograms (Figure 5). The only exception to this was the stochastic character maps for the locus-inferred phylogram, which estimated significantly lower total rates and only 7 total transitions across the tree. The true ancestral character history is almost always unknown, and so here we were more interested in precision rather than accuracy. As such, we found that the estimated evolutionary parameters were more consistent between SNP and locus chronograms compared to phylograms (Figure 5). This likely reflects the drastic differences in absolute branch lengths between SNP and locus phylograms (Figure 2A), which is eliminated in chronograms.

**Figure 6.**
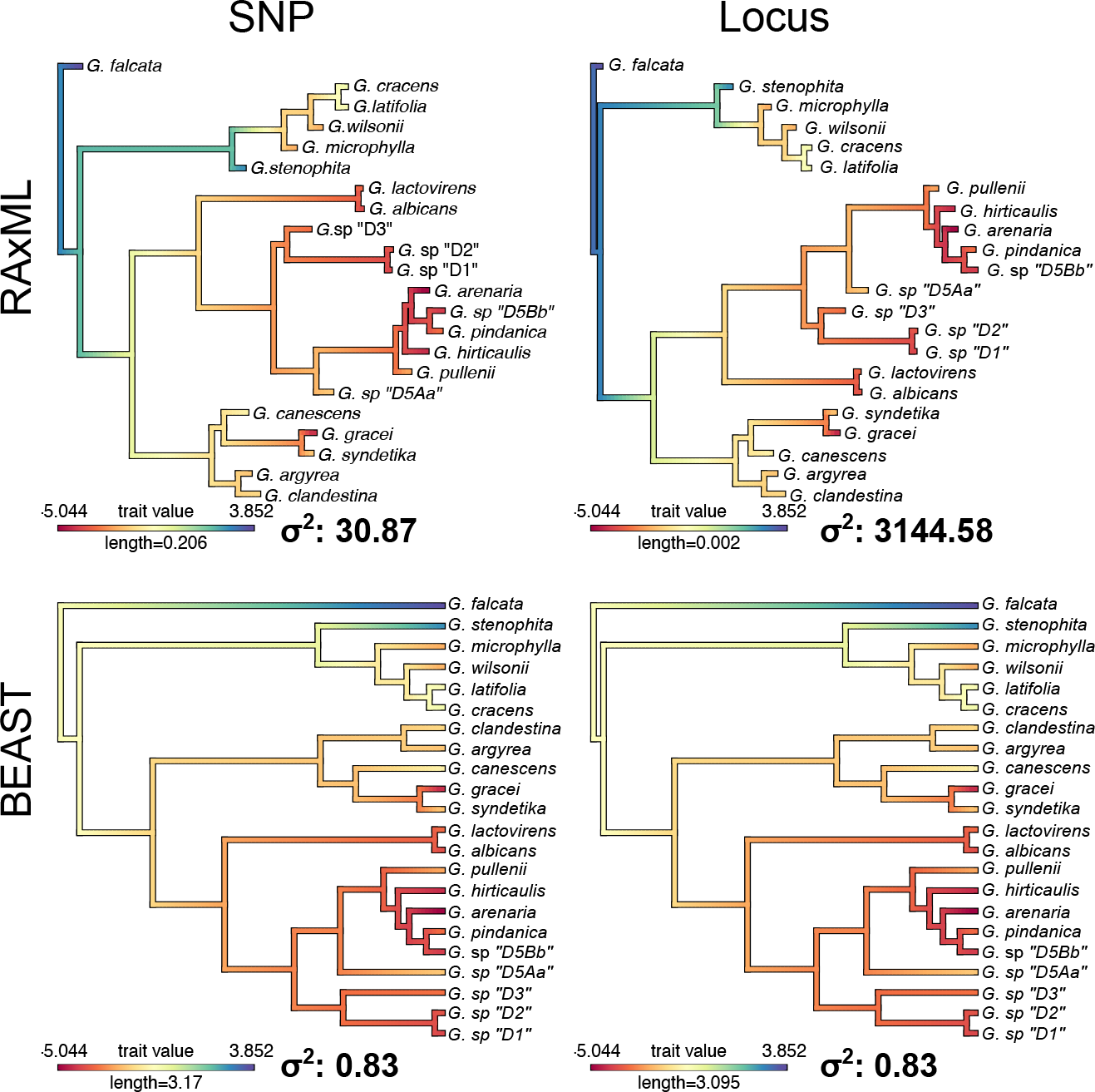
Summarized results of ancestral character estimation of continuous character states for two representative phylograms and chronograms from the 45% filtering stringency. Branches are colored by their estimated continuous state based on the associated legend. The σ2 represents the estimated Brownian motion parameter. The data were modeled with a σ2 equal to 1.

These downstream results are hardly surprising given that rates of morphological change in phylograms are denoted as the number of morphological changes per nucleotide substitution per site. This rate does not make sense biologically, unless the tree was inferred with genes that underlie the morphological traits of interest and if mutations in those genes are known to directly correlate with particular shifts in phenotype (Litsios and Salamin, 2012). Given the drastic differences and sensitivity of modeled parameters to branch lengths, we recommend authors use chronograms when conducting phylogenetic comparative methods.

### Recommendations

There are many considerations that should be taken into account when determining which sequencing method is most appropriate. For instance, the number of taxa, taxon sampling, crown age, heterozygosity, and the biological question of interest are all important to consider before any data are generated. If researchers conclude that a SNP-based approach is best, the results from our analyses suggest the following recommendations: **1**. Researchers can use either SNPs or loci in their analyses to produce reliable inferences. This is supported by the equivalent results in our phylogenomic inference, including overall topology, divergence time estimation, and PCMs on chronograms. **2**. Use moderate filtering (∼30-45%) in SNP retention as over-filtering can lead to inaccurate branch lengths and in some cases minor topological differences. On the other hand, under-filtering SNPs increases computation time with minimal benefit to phylogenetic inference. **3**. Use chronograms in downstream phylogenetic comparative methods as phylograms are highly sensitive to differences in branch lengths leading to imprecise evolutionary metrics. We recognize that many of these decisions depend on the questions researchers hope to answer, which should drive the methods they use. However, we hope our framework can be helpful for phylogenetic analyses in the genomics age and can influence some of the methodological decisions taken by future researchers.

## Author contributions

GYD, LG, ADG, CJ, HRP, DW, JSS, and JBL conceptually designed the project.

JSS and JBL conducted the analyses and drafted the manuscript.

All authors contributed to editing the final draft of the manuscript.

## Acknowledgements

This manuscript came about after a semester-long discussion group at Cornell centered around phylogenomics. Would like to thank others that made the discussion group fruitful, including Yanã Campos Rizzieri, Israel Cunha Neto, Caroline Dowling, Joyce Onyenedum, Mônica Ulysséa, and Shirley Zhang. We also thank Sue Sherman-Broyles for generating the GBS data that was used in this study. This work was partially funded by the US Department of Agriculture National Institute of Food and Agriculture Hatch projects 1014310 and 7002754 (JJD) and the National Science Foundation 2208845 (JSS).

## Data availability statement

Raw sequencing reads are available on the Sequence Read Archive with the accession numbers presented in Table 1. Analyses and plotting scripts are available in Supplemental Materials.

