## Supporting Information for "Comparative phylogenomic analyses of SNP versus full locus datasets: insights and recommendations for researchers"

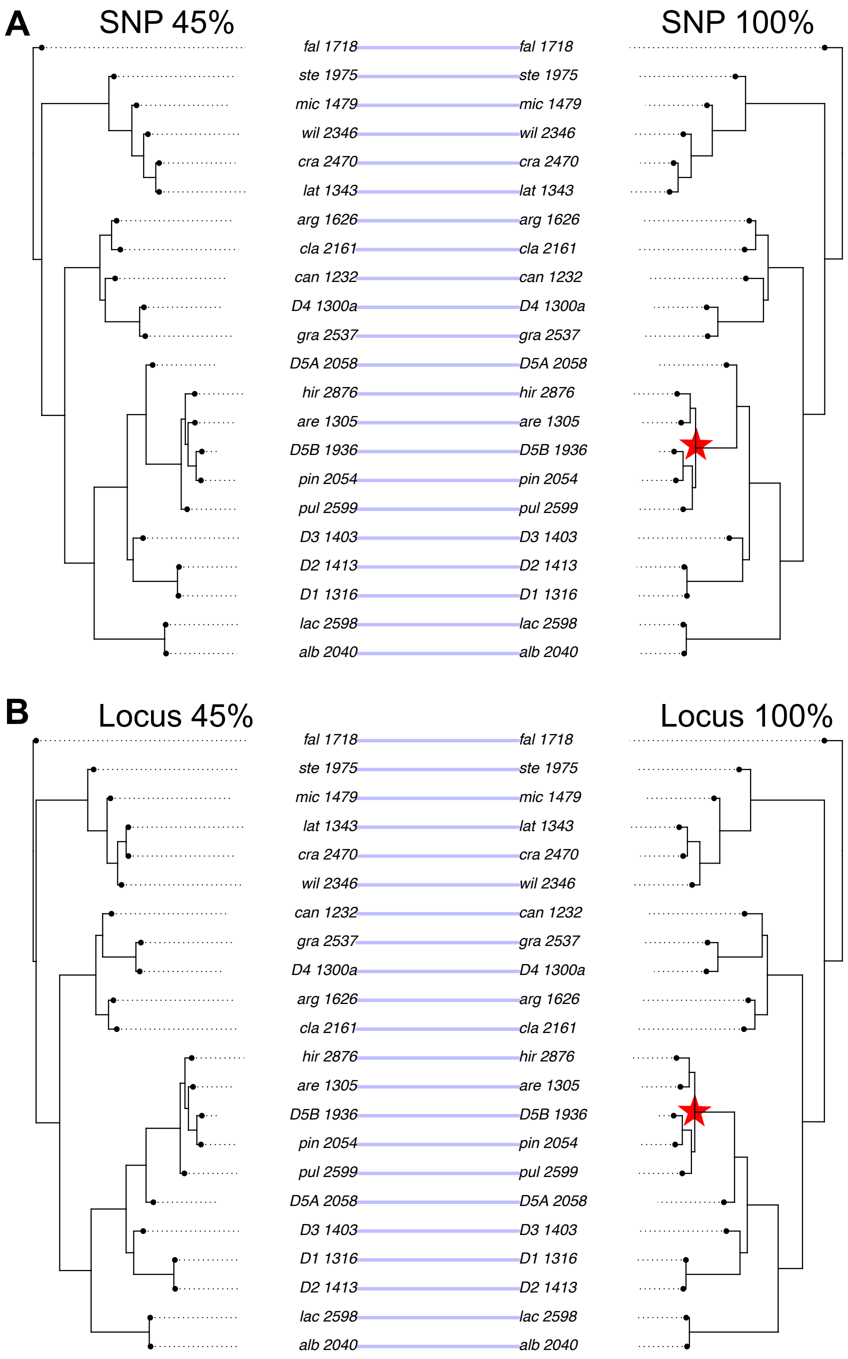

Figure S1. Depiction of the one topological incongruence between 100% filtering stringency for A) SNP and B) locus data. Star represents topological incongruence. The tree depicted with 45% filtering represents the overall topology across all other filtering stringencies and data types.

Supplemental Table 1. Amount of missing data for each accession in each dataset as calculated by VCFtools in the resulting VCF file.

| Accession | 100% - Variant | 100% - locus | 90% - Variant | 90% - Locus | 75% - Variant | 75% - Locus | 60% - Variant | 60% - Locus | 45% - Variant | 45% - Locus | 30% - Variant | 30% - Locus | 15% - Variant | 15% - Locus | 0% - Variant | 0% - Locus |
|---|---|---|---|---|---|---|---|---|---|---|---|---|---|---|---|---|
| Glycine albicans | 0 | 0.115 | 0.029 | 0.139 | 0.103 | 0.208 | 0.163 | 0.261 | 0.266 | 0.361 | 0.370 | 0.488 | 0.474 | 0.692 | 0.479 | 0.878 |
| Glycine arenaria | 0 | 0.116 | 0.056 | 0.158 | 0.160 | 0.247 | 0.265 | 0.337 | 0.404 | 0.464 | 0.502 | 0.563 | 0.600 | 0.704 | 0.604 | 0.943 |
| Glycine argyrea | 0 | 0.115 | 0.023 | 0.130 | 0.061 | 0.159 | 0.111 | 0.202 | 0.209 | 0.291 | 0.311 | 0.414 | 0.422 | 0.611 | 0.428 | 0.904 |
| Glycine canescens | 0 | 0.015 | 0.032 | 0.042 | 0.083 | 0.087 | 0.141 | 0.151 | 0.266 | 0.292 | 0.376 | 0.436 | 0.472 | 0.622 | 0.478 | 0.900 |
| Glycine clandestina | 0 | 0.115 | 0.026 | 0.134 | 0.063 | 0.162 | 0.115 | 0.203 | 0.200 | 0.283 | 0.300 | 0.405 | 0.411 | 0.601 | 0.417 | 0.889 |
| Glycine sp "cracens" | 0 | 0.177 | 0.036 | 0.200 | 0.154 | 0.285 | 0.287 | 0.405 | 0.429 | 0.536 | 0.533 | 0.647 | 0.608 | 0.704 | 0.610 | 0.907 |
| Glycine tomentella D1 | 0 | 0.014 | 0.014 | 0.026 | 0.057 | 0.065 | 0.104 | 0.111 | 0.216 | 0.233 | 0.313 | 0.341 | 0.406 | 0.564 | 0.411 | 0.871 |
| Glycine tomentella D2 | 0 | 0.013 | 0.042 | 0.051 | 0.109 | 0.112 | 0.160 | 0.166 | 0.272 | 0.289 | 0.366 | 0.394 | 0.457 | 0.608 | 0.462 | 0.875 |
| Glycine tomentella D3 | 0 | 0.015 | 0.035 | 0.044 | 0.082 | 0.086 | 0.129 | 0.132 | 0.246 | 0.257 | 0.346 | 0.367 | 0.442 | 0.581 | 0.448 | 0.872 |
| Glycine syndetika | 0 | 0.015 | 0.035 | 0.047 | 0.090 | 0.099 | 0.150 | 0.163 | 0.266 | 0.294 | 0.370 | 0.428 | 0.457 | 0.600 | 0.462 | 0.865 |
| Glycine tomentella D5A | 0 | 0.116 | 0.021 | 0.128 | 0.057 | 0.156 | 0.105 | 0.194 | 0.224 | 0.308 | 0.337 | 0.426 | 0.455 | 0.633 | 0.460 | 0.902 |
| Glycine tomentella D5B | 0 | 0.115 | 0.021 | 0.130 | 0.064 | 0.165 | 0.139 | 0.229 | 0.265 | 0.344 | 0.369 | 0.451 | 0.482 | 0.622 | 0.488 | 0.917 |
| Glycine falcata | 0 | 0.119 | 0.373 | 0.454 | 0.526 | 0.580 | 0.622 | 0.669 | 0.713 | 0.760 | 0.773 | 0.828 | 0.821 | 0.903 | 0.823 | 0.957 |
| Glycine gracei | 0 | 0.116 | 0.025 | 0.134 | 0.076 | 0.176 | 0.144 | 0.235 | 0.247 | 0.336 | 0.353 | 0.467 | 0.461 | 0.652 | 0.466 | 0.905 |
| Glycine hirticaulis | 0 | 0.105 | 0.048 | 0.144 | 0.132 | 0.218 | 0.227 | 0.301 | 0.368 | 0.430 | 0.468 | 0.531 | 0.569 | 0.686 | 0.574 | 0.932 |
| Glycine lactovirens | 0 | 0.115 | 0.031 | 0.137 | 0.105 | 0.206 | 0.166 | 0.260 | 0.273 | 0.365 | 0.381 | 0.499 | 0.483 | 0.703 | 0.488 | 0.891 |
| Glycine latifolia | 0 | 0.178 | 0.037 | 0.205 | 0.172 | 0.301 | 0.310 | 0.426 | 0.458 | 0.567 | 0.564 | 0.681 | 0.636 | 0.731 | 0.639 | 0.921 |
| Glycine microphylla | 0 | 0.168 | 0.036 | 0.192 | 0.145 | 0.273 | 0.278 | 0.396 | 0.427 | 0.540 | 0.537 | 0.659 | 0.614 | 0.718 | 0.618 | 0.929 |
| Glycine pindanica | 0 | 0.105 | 0.021 | 0.120 | 0.063 | 0.154 | 0.130 | 0.213 | 0.277 | 0.356 | 0.395 | 0.479 | 0.507 | 0.646 | 0.512 | 0.920 |
| Glycine pullenii | 0 | 0.116 | 0.080 | 0.186 | 0.193 | 0.283 | 0.291 | 0.364 | 0.413 | 0.473 | 0.500 | 0.561 | 0.594 | 0.702 | 0.598 | 0.941 |
| Glycine stenophita | 0 | 0.116 | 0.043 | 0.150 | 0.095 | 0.196 | 0.199 | 0.304 | 0.334 | 0.442 | 0.441 | 0.568 | 0.534 | 0.685 | 0.539 | 0.918 |
| Glycine sp "wilsonii" | 0 | 0.114 | 0.026 | 0.129 | 0.079 | 0.181 | 0.186 | 0.293 | 0.303 | 0.409 | 0.403 | 0.526 | 0.492 | 0.621 | 0.496 | 0.891 |

Supplemental Table 2. Wilcoxon signed rank pairwise comparisons of branch length and node support for the different data types and filtering strategies.

| Data_type | Trait | Filtering_level | | | | | | | |
|---|---|---|---|---|---|---|---|---|---|
| | | | 0 | 100 | 15 | 30 | 45 | 60 | 75 |
| SNP | Node support | 100 | 1 | - | - | - | - | - | - |
| SNP | Node support | 15 | 1 | 1 | - | - | - | - | - |
| SNP | Node support | 30 | 1 | 0.069 | 1 | - | - | - | - |
| SNP | Node support | 45 | 1 | 0.069 | 1 | 1 | - | - | - |
| SNP | Node support | 60 | 1 | 1 | 1 | 1 | 1 | - | - |
| SNP | Node support | 75 | 1 | 1 | 1 | 1 | 1 | 1 | - |
| SNP | Node support | 90 | 1 | 1 | 1 | 1 | 1 | 1 | 1 |
| SNP | Branch length | 100 | 0.022 | - | - | - | - | - | - |
| SNP | Branch length | 15 | 0.777 | 0.031 | - | - | - | - | - |
| SNP | Branch length | 30 | 1 | 0.021 | 1 | - | - | - | - |
| SNP | Branch length | 45 | 0.683 | 0.089 | 0.335 | 1 | - | - | - |
| SNP | Branch length | 60 | 1 | 0.054 | 1 | 1 | 1 | - | - |
| SNP | Branch length | 75 | 0.012 | 0.729 | 0.127 | 0.91 | 1 | 1 | - |
| SNP | Branch length | 90 | 0.525 | 1 | 0.939 | 0.29 | 1 | 1 | 1 |
| Locus | Node support | 100 | 0.86 | - | - | - | - | - | - |
| Locus | Node support | 15 | 1 | 0.86 | - | - | - | - | - |
| Locus | Node support | 30 | 1 | 0.16 | 1 | - | - | - | - |
| Locus | Node support | 45 | 1 | 0.16 | 1 | 1 | - | - | - |
| Locus | Node support | 60 | 1 | 1 | 1 | 1 | 1 | - | - |
| Locus | Node support | 75 | 1 | 0.16 | 1 | 1 | 1 | 1 | - |
| Locus | Node support | 90 | 1 | 1 | 1 | 1 | 1 | 1 | 1 |
| Locus | Node support | | | | | | | | |
| Locus | Branch length | 100 | 9.80E-07 | - | - | - | - | - | - |
| Locus | Branch length | 15 | 2.30E-05 | 1 | - | - | - | - | - |
| Locus | Branch length | 30 | 9.10E-07 | 1 | 1 | - | - | - | - |
| Locus | Branch length | 45 | 4.80E-07 | 0.70587 | 0.13203 | 0.06386 | - | - | - |
| Locus | Branch length | 60 | 5.10E-07 | 0.00042 | 0.00132 | 1 | 1 | - | - |
| Locus | Branch length | 75 | 9.50E-07 | 0.81516 | 0.02758 | 1 | 1 | 1 | - |
| Locus | Branch length | 90 | 8.50E-07 | 1 | 0.52462 | 1 | 1 | 1 | 1 |
